# Fragle: Universal ctDNA quantification using deep learning of fragmentomic profiles

**DOI:** 10.1101/2023.07.28.550922

**Authors:** Guanhua Zhu, Chowdhury Rafeed Rahman, Victor Getty, Denis Odinokov, Probhonjon Baruah, Hanaé Carrié, Avril Joy Lim, Yu Amanda Guo, Zhong Wee Poh, Ngak Leng Sim, Ahmed Abdelmoneim, Yutong Cai, Lakshmi Lakshmanan, Danliang Ho, Saranya Thangaraju, Polly Poon, Yi Ting Lau, Anna Gan, Sarah Ng, Si-Lin Koo, Dawn Q. Chong, Brenda Tay, Tira J. Tan, Yoon Sim Yap, Aik Yong Chok, Matthew Chau Hsien Ng, Patrick Tan, Daniel Tan, Limsoon Wong, Pui Mun Wong, Iain Beehuat Tan, Anders Jacobsen Skanderup

## Abstract

Quantification of circulating tumor DNA (ctDNA) levels in blood enables non-invasive surveillance of cancer progression. Fragle is an ultra-fast deep learning-based method for ctDNA quantification directly from cell-free DNA fragment length profiles. We developed Fragle using low-pass whole genome sequence (lpWGS) data from multiple cancer types and healthy control cohorts, demonstrating high accuracy, and improved lower limit of detection in independent cohorts as compared to existing tumor-naïve methods. Uniquely, Fragle is also compatible with targeted sequencing data, exhibiting high accuracy across both research and commercial targeted gene panels. We used this method to study longitudinal plasma samples from colorectal cancer patients, identifying strong concordance of ctDNA dynamics and treatment response. Furthermore, prediction of minimal residual disease in resected lung cancer patients demonstrated significant risk stratification beyond a tumor-naïve gene panel. Overall, Fragle is a versatile, fast, and accurate method for ctDNA quantification with potential for broad clinical utility.

## Introduction

The death of non-malignant cells, primarily of the hematopoietic lineage, releases cell-free DNA (cfDNA) into the blood circulation ^1^. In cancer patients, the blood plasma also carries circulating tumor DNA (ctDNA), enabling non-invasive diagnostics and disease surveillance ^2^. The ability to monitor tumor growth dynamics based on ctDNA levels in the blood provides a promising non-invasive approach to track disease progression during therapy and clinical trials ^3–5^.

Ultra-deep targeted cfDNA sequencing assays are often preferred in the clinic due to their ability to identify actionable mutations. While mutation variant allele frequencies (VAFs) can be used to approximate ctDNA levels, not all tumors will have mutations covered by a given targeted sequencing gene panel. Furthermore, the accuracy of this approximation depends on sample-specific and treatment-dynamic properties such as mutation clonality, copy number, as well as potential confounding noise from clonal hematopoiesis ^6^. Existing methods developed for ctDNA quantification are not directly compatible with targeted sequencing panels. These methods require either low-pass whole genome sequencing (lpWGS) data ^7^, DNA methylation profiling ^8, 9^, or modifications to the targeted sequencing panel ^10^. Thus, there is an unmet need to develop accurate and orthogonal approaches for ctDNA quantification that can generalize across patients, tumor types, and sequencing modalities.

The fragment length distribution of cfDNA in plasma has a mode of ∼166 base pairs (bp) as nucleosome-bound cfDNA molecules display increased protection from DNA degradation ^11^. cfDNA fragments from cancer patients tend to be shorter than those from healthy individuals, typically with a higher proportion of fragments under 150bp ^12–14^. Shorter cfDNA fragments have also been observed in plasma bisulfite sequencing data from cancer patients ^15^. The size profile of these shorter fragments from cancer patients also exhibits increased 10-bp oscillation amplitude in the range 90-145bp ^16^. cfDNA from cancer patients may also display a higher proportion of fragments longer than 180bp ^12, 16^. Other studies have indicated that variation in fragment lengths in cancer patients could be position-dependent within the genome ^17^. These observations have motivated studies exploring how cfDNA fragment length properties can be used to classify cfDNA samples from cancer patients and healthy individuals ^12, 15–23^. Here, we developed *Fragle*, a multi-stage machine learning model that quantifies ctDNA levels from a cfDNA fragment length density distribution. Using an in-silico data augmentation approach, we trained and evaluated Fragle on ∼4000 lpWGS samples across distinct cancer types and healthy cohorts. We evaluated the accuracy and the lower limit of detection (LoD) in independent cohorts and cancer types. Intriguingly, we demonstrate that Fragle can also be applied to cfDNA fragmentomic profiles obtained from targeted sequencing panels. Using this feature, we applied Fragle to longitudinal plasma samples to explore the correlation of ctDNA dynamics and treatment response measured through radiographic imaging. Finally, to explore the use of Fragle for detection of minimal residual disease (MRD), we analyzed ctDNA levels in a cohort of 162 resected lung cancer patients with plasma profiled using a commercial targeted sequencing panel at the landmark timepoint (∼30 days following surgery).

## Results

### Quantitative prediction of ctDNA levels from fragmentomic data

We assembled a discovery cohort comprising lpWGS data from 325 cancer plasma samples from 4 cancer types (colon, breast, liver, and ovarian cancer) and 101 plasma samples from healthy individuals (Fig. 1, Suppl. Data 1). In this dataset, we estimated ground-truth ctDNA levels using multiple methods (see Methods, Suppl. Data 2), and the cancer samples were further selected based on ctDNA levels (≥3%, N = 164, Suppl. Fig. 1). Using a large-scale data augmentation approach, we performed in-silico dilution of these cancer samples and the 101 healthy control samples, generating ∼4000 mixture samples with variable ctDNA fractions for model training (Methods, Fig. 1, Suppl. Data 3). To explore how cfDNA fragment length distributions could predict ctDNA levels in a sample, we derived raw fragment length density distributions using paired-end reads in each sample. Raw density distributions were further normalized and transformed, revealing local differences in the fragment length distributions associated with ctDNA levels in the samples (Fig. 1, see Methods). The transformed fragment length distributions, in combination with their labels in the form of ground-truth ctDNA levels, served as input to a multi-stage supervised machine learning approach. We employed two parallel sub-models, each designed for either low- or high-ctDNA fraction samples, followed by a model that selects the final predicted ctDNA fraction from the output of the two sub-models (see Methods). The two sub-models performed well for the intended low- and high-ctDNA samples, respectively (Suppl. Fig. 2), while the final combined model achieved the lowest overall prediction error (MAE = 3.2%) as compared to individual sub-models (MAE: 4.0% and 3.3% for low- and high-ctDNA sub-models). Notably, although the improvement in overall MAE is modest compared to the high-ctDNA sub-model, the final combined model significantly improved the prediction accuracy for healthy samples (MAE: 0.5% vs. 1.0%) and specificity at an LoD of 1% (86% vs 68%).

**Fig. 1.**
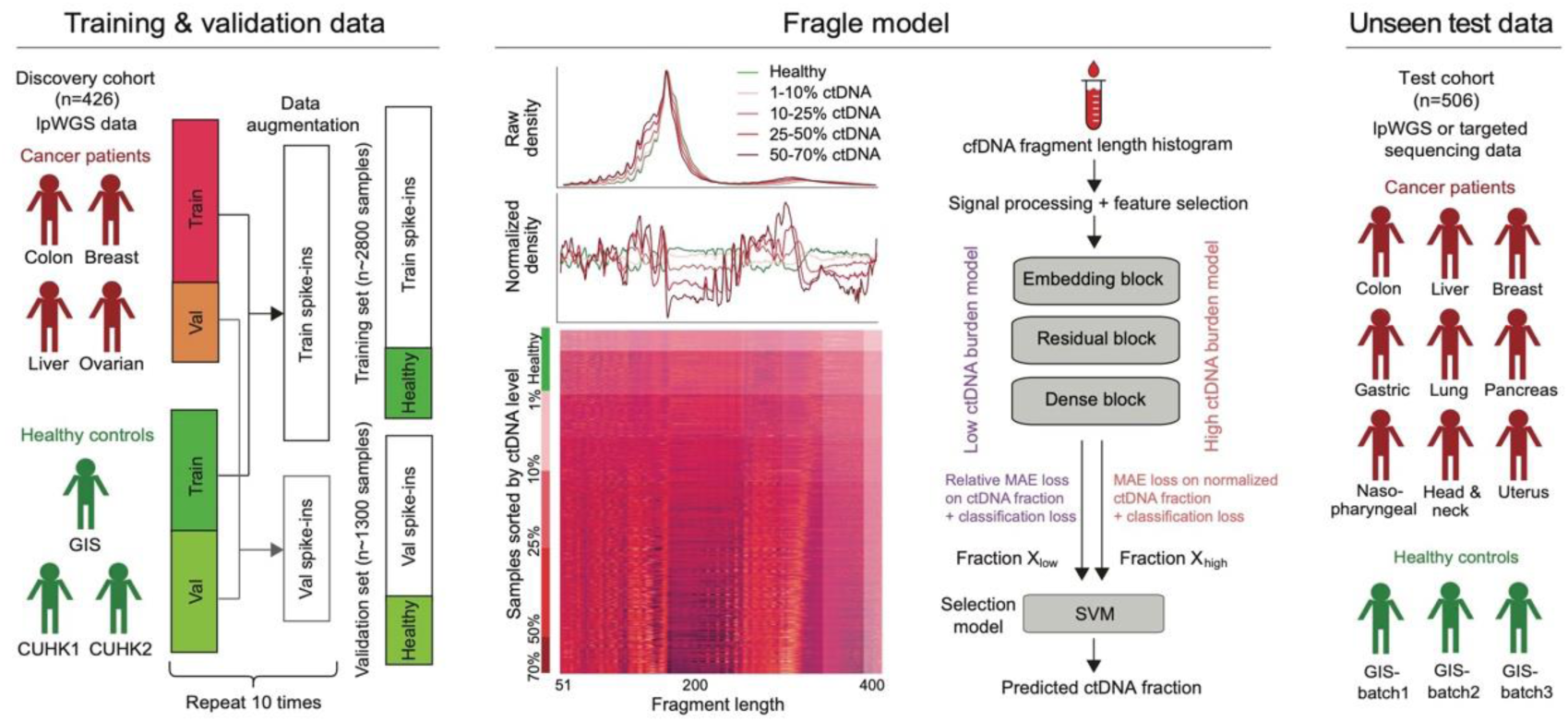
Overview of Fragle. Fragle is a multi-stage machine learning-based model that estimates the ctDNA level in a blood sample from the cfDNA fragment length density distribution. Fragle was trained using a large-scale data augmentation and cross-validation approach and was further tested using unseen samples from multiple cancer types and healthy control cohorts.

The model was trained and evaluated using cross-validation, demonstrating high predictive accuracy on validation samples across all 4 cancer types (Fig. 2a-d, Suppl. Data 4): Colorectal (mean absolute error (MAE) = 3.3%; Pearson *r* = 0.92), breast (MAE = 3.6%; *r* = 0.94), liver (MAE = 3.1%; *r* = 0.81), and ovarian cancer (MAE = 3.9%; *r* = 0.67). The lower concordance for ovarian cancer could be attributed to samples from one patient; removal of these samples increased the correlation to *r* = 0.88 (Suppl. Fig. 3).

**Fig. 2.**
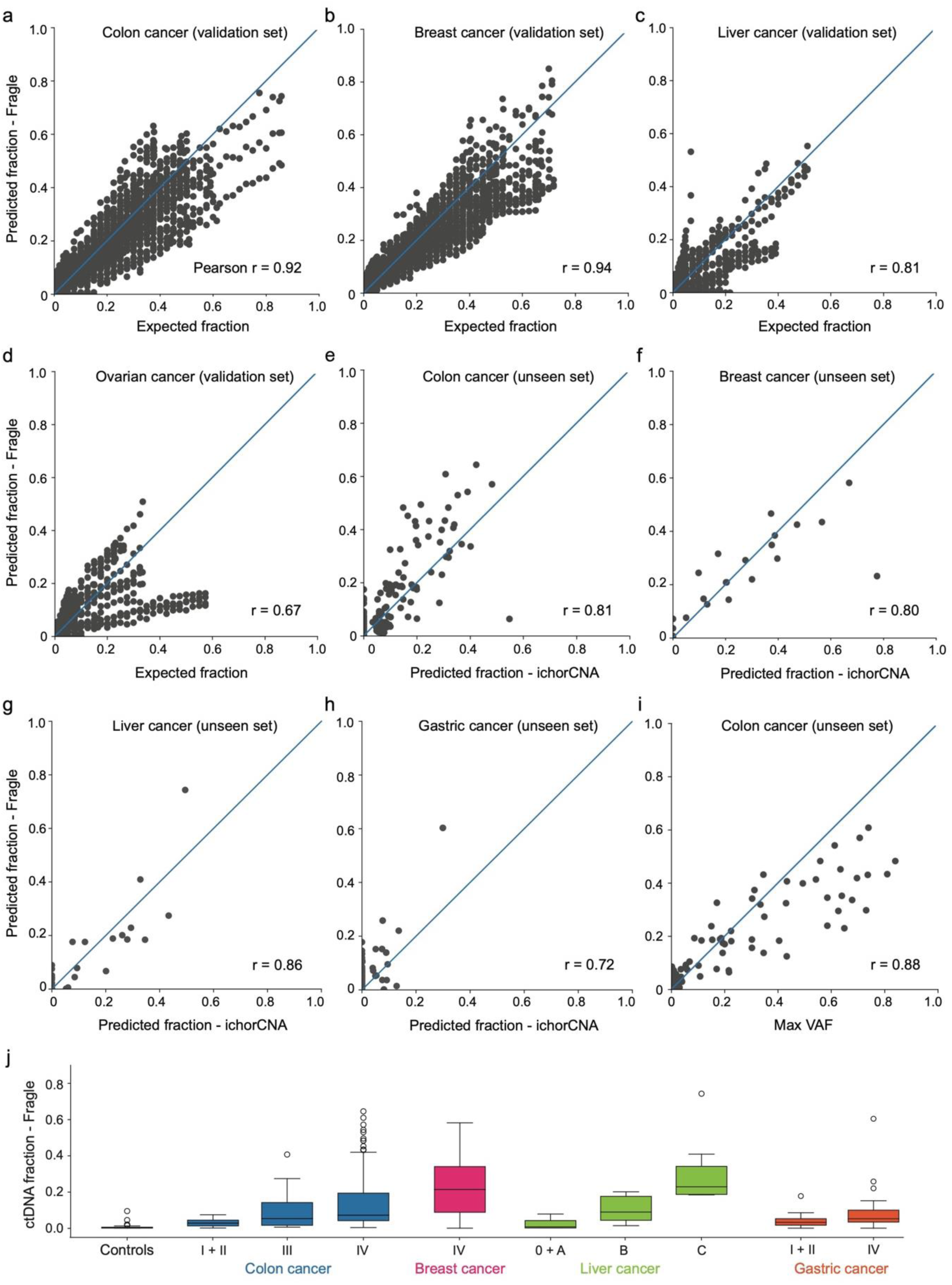
ctDNA quantification in validation and unseen cohorts. **(a-d)** Comparison between expected and predicted ctDNA levels for colorectal (CRC), breast (BRCA), liver (HCC), and ovarian (OV) cancer samples in the validation sets. (**e-h)** Comparing ichorCNA and Fragle predicted ctDNA levels in unseen samples from colorectal (N = 172), breast (N = 23), liver (N = 34), and gastric cancer patients (N = 74). **i)** Colorectal cancer plasma samples subjected to both lpWGS and targeted sequencing; comparison of Fragle predicted ctDNA levels (lpWGS) and maximum VAFs (N = 86; samples with detectable somatic mutations). j) Predicted ctDNA levels in plasma samples from cancer patients grouped according to tumor stages.

We trained the final Fragle model on the full discovery cohort (see Methods) and tested its performance on additional cohorts of unseen plasma lpWGS samples. We observed a strong correlation between Fragle and ichorCNA-based ctDNA fraction estimates across unseen cohorts of colorectal cancer (*r* = 0.81; *P* = 6.8e-41; N = 172; Fig. 2e), breast cancer (*r* = 0.80; *P* = 3.7e-06; N = 23; Fig. 2f), liver cancer (*r* = 0.86; *P* = 5.1e-10; N = 34; Fig. 2g), and gastric cancer (*r* = 0.72; *P* = 3.4e-13; N = 74; Fig. 2h). We also tested Fragle on a mixed cohort of cancer types not included in the discovery set, including lung, nasopharyngeal, as well as head and neck cancers (r = 0.63, 0.75, and 0.23; combined *P* = 1.3e-4, n = 10 for each cancer type; Suppl. Fig. 4). In the unseen colorectal cancer cohort, we also performed targeted gene sequencing and identified high-confidence somatic mutations in 86 samples (Suppl. Data 5, see Methods). These data demonstrated high concordance between mutation VAFs and Fragle-predicted ctDNA levels (*r* = 0.88; *P* = 3.8e-28; Fig. 2i). Expectedly, higher ctDNA fractions were generally observed in the patients with late-stage tumors (Fig. 2j; Suppl. Data 6). Furthermore, we observed a significant difference between ctDNA levels estimated for early-stage cancers (stages 1 and 2; colon, liver, and gastric cancer) and healthy controls (*P* = 1.3e-9, Wilcoxon rank sum test).

We trained Fragle using samples each comprising 10 million cfDNA fragments, equivalent to ∼1x WGS using 151bp paired-end sequencing. To further evaluate the sequencing coverage requirements for Fragle, we down-sampled WGS samples from the unseen test cohort to render samples with fewer fragments, ranging from 5 million (0.5x) to as low as 10 thousand fragments (0.001x). At 500K fragments (0.05x), Fragle demonstrated excellent concordance (*r* = 0.97) with the predictions from the original 1x WGS samples (Suppl. Fig. 5). The correlation was maintained when further down-sampling to 250K fragments, but with some discrepancies observed for some low ctDNA fraction samples (Suppl. Fig. 5). These results suggest that whole-genome coverage of ∼500K (0.05x) fragments provides a good trade-off between prediction accuracy and sequencing cost. In addition, we tested the computational requirements of Fragle as a software tool. Fragle processed a 1x-coverage WGS sample in ∼50 seconds using a single processor and required low memory usage independent of the sample sequencing depth (Suppl. Fig. 6).

### Determination of the lower limit of detection

To explore the lower limit of detection (LoD) for the model, we first observed that Fragle predicted very low ctDNA fractions (median = 0.07%) for the healthy samples in the validation sets. In this healthy cohort, Fragle demonstrated 86% specificity at a 1% LoD level, increasing to 95% at 3% LoD (Suppl. Data 7). Furthermore, the model could differentiate between healthy and low-ctDNA level samples at the 1% ctDNA level (Wilcoxon rank sum test *P* = 2.5e-24, Fig. 3a), indicating a ∼1% LoD in these samples. Similarly, we examined the performance of Fragle for classification of healthy and cancer samples in the validation sets. Using cancer samples with a ctDNA level ≥1% in the validation sets, Fragle demonstrated an area under the curve (AUC) of 0.93 (Fig. 3b), higher than ichorCNA (AUC = 0.88) applied to the same samples. Notably, after limiting the analysis to the samples in which the ground-truth ctDNA fraction was estimated from a consensus of multiple methods, Fragle further outperformed ichorCNA in classifying cancer and healthy samples (AUC: 0.98 vs. 0.92; Suppl. Fig. 7). Expectedly, the AUC increased further when filtering out low-ctDNA burden samples (Suppl. Fig. 8). Fragle and ichorCNA achieved AUCs of 0.97 and 0.94, respectively, when excluding cancer samples with ctDNA levels below ichorCNA’s LoD of 3% (Suppl. Fig. 9). As an additional comparison, we explored other fragment length features previously used for the classification of cancer and healthy samples ^16^, and trained a random forest model on the discovery cohort using 4 features derived from the fragment length distribution (see Methods). This 4-feature model demonstrated substantially lower classification accuracy (AUC = 0.79) than Fragle in the validation cohort.

**Fig. 3.**
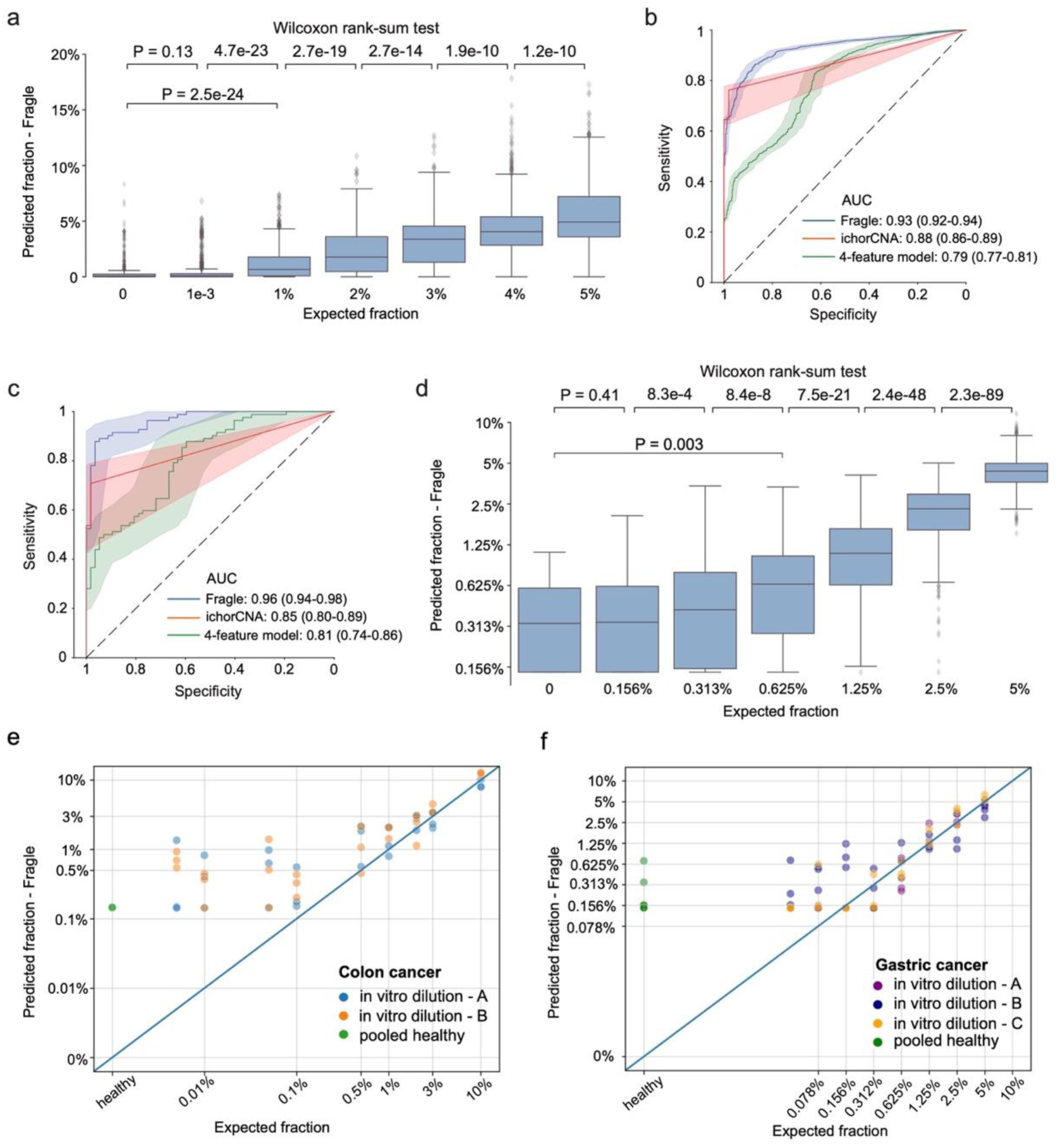
Lower limit of detection. **a)** Predicted ctDNA fractions for healthy and low-ctDNA level samples in validation set samples. Boxplots are represented by median and interquartile range (IQR), with +/–1.5 IQR as whiskers. **b)** ROC analyses for classification of healthy control and cancer (≥1% ctDNA) samples (validation samples). AUC values with 95% confidence intervals are shown. **c)** ROC analysis for classification of cancer (N=86) and healthy (N=57, 3 distinct cohorts) plasma samples in the unseen test cohort. AUC values with 95% confidence intervals are shown. **d)** Predicted ctDNA fractions for healthy and low-ctDNA level samples using in silico dilution of 20 cancer samples (unseen cohort). Boxplots are represented by median and interquartile range (IQR), with +/–1.5 IQR as whiskers. **e)** Expected vs. predicted ctDNA fractions using in vitro ctDNA dilution for 2 colorectal cancer samples. **f)** Expected vs. predicted ctDNA fractions using in vitro ctDNA dilution for 3 gastric cancer samples.

We further evaluated the LoD using unseen test samples. We used cfDNA samples from CRC patients with detectable mutations as positive cancer samples (N = 86, Suppl. Data 8) and all healthy plasma samples from the unseen cohorts as negatives (N = 57, Suppl. Data 9). Fragle demonstrated an AUC of 0.96 using these samples, outperforming the other models on the same set of samples (ichorCNA = 0.85, 4-feature model = 0.81; Fig. 3c; Suppl. Data 10). These results were further confirmed using an in-silico dilution experiment. This experiment involved 13 unseen colon cancer and 7 unseen breast cancer samples with high ctDNA burden (>10%), concordantly estimated by Fragle and ichorCNA (see Methods, Suppl. Data 11). In this dilution experiment, Fragle could differentiate healthy from low-ctDNA samples down to the 0.5-1% ctDNA level (*P* = 0.003, healthy vs. 0.625% ctDNA fraction samples, Fig. 3d; Suppl. Fig. 10).

To further examine these results using physical samples, we performed similar dilution experiments in vitro. The first experiment comprised serial dilutions of two high-ctDNA level CRC plasma samples, with samples progressively diluted using pooled cfDNA from healthy individuals (see Methods). Across 3 technical replicates, Fragle accurately predicted ctDNA fractions for both patients down to ∼1% ctDNA level, with healthy samples consistently predicted <1% ctDNA (Fig. 3e). For low-ctDNA samples with 1-3% diluted ctDNA fraction, the detection rate was 94% at an LoD of 1%, outperforming ichorCNA with a detection rate of 67% (Suppl. Data 12). The second experiment comprised in vitro serial dilutions of 3 high-ctDNA plasma samples from gastric cancer patients (each with 3 technical replicates, see Methods). The results from this experiment mirrored our previous observations, with the method accurately quantifying ctDNA down to the 0.5-1% level and predicting <1% ctDNA for healthy samples (Fig. 3f, Suppl. Data 13). Overall, these results collectively suggest that Fragle can quantify and detect ctDNA with an LoD of ∼1%.

### Application of Fragle to targeted sequencing data

Targeted gene sequencing of plasma samples is routinely used for tumor genotyping in the clinic. However, absolute ctDNA quantification based on mutation VAFs remains challenging using targeted sequencing. For example, samples may not have clonal mutations covered by the panel, and non-cancer variants associated with clonal hematopoiesis could introduce noise ^24^. To explore whether Fragle could quantify ctDNA levels using targeted sequencing data, we analyzed four cfDNA cohorts having both lpWGS and targeted sequencing data (Fig. 4a, see Methods). Using standard on-target reads obtained from the targeted sequencing data, Fragle tended to overestimate the ctDNA burden as compared to the lpWGS data (Fig. 4b). We then evaluated the method on off-target reads, which are often filtered and ignored in a targeted sequencing experiment. Remarkably, we observed strong concordance of predictions based on lpWGS and off-target reads across all four cohorts: breast cancer samples from the discovery cohort (r = 0.86, *P* = 0.001, N = 10), colon cancer samples from the discovery cohort (r = 0.96, *P* = 1.64e-30, N = 56), colon cancer samples from the unseen cohort (r = 0.97, *P* = 3.69e-58, N = 109), and metastatic gastric cancer samples from the unseen cohort (r = 0.96, *P* = 2.9e-27, N = 49) (Fig. 4c). We found that the targeted sequencing samples contained between 100K to 10M off-target fragments (equivalent to ∼0.01-1.0X WGS) across the different samples, with >95% of samples having >250K off-target fragments (∼0.025X; Suppl. Fig. 11). Expectedly, the off-target coverage levels showed a linear relationship to on-target coverage across samples (Suppl. Fig. 11). To further explore if these results generalize to other targeted sequencing assays, we evaluated a cohort of 116 plasma samples subjected to a liquid biopsy gene panel from a commercial vendor (Foundation Medicine) ^25^. Since these samples did not have matched lpWGS data, we approximated ctDNA levels using the maximum VAFs reported by the company after filtering out germline variants (Suppl. Data 14; Methods). In this cohort comprising samples from 5 different cancer types, we observed that ctDNA levels estimated from off-target reads were generally concordant with the reported VAFs (r = 0.62, *P* = 1.4e-13, N = 116; Fig. 4d). Overall, these results support that Fragle can estimate ctDNA levels using both lpWGS and targeted sequencing data.

**Fig. 4.**
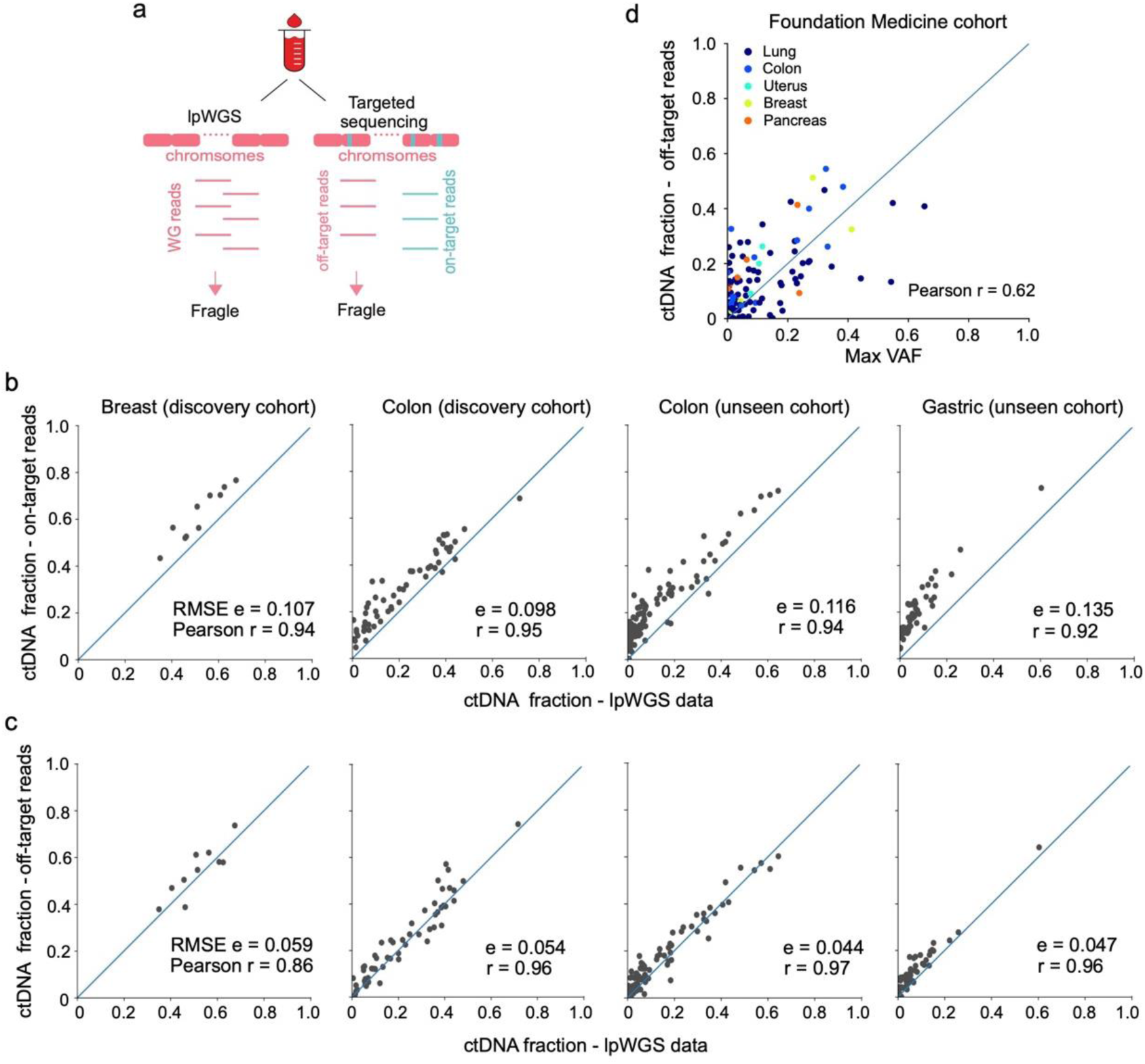
Application of Fragle to targeted sequencing data. **a)** Application of Fragle to samples having both lpWGS and targeted gene panel sequencing data. **b)** ctDNA levels predicted using lpWGS data and on-target reads from targeted sequencing samples. **c)** ctDNA levels predicted using lpWGS data and off-target reads from targeted sequencing samples. **d)** Targeted sequencing data generated with commercial liquid biopsy assay (Foundation Medicine, N=116). Correlation of maximum VAFs (reported by the company, germline variants filtered) and Fragle-predicted ctDNA levels using off-target reads.

### Tracking ctDNA dynamics and disease progression from targeted sequencing

Having demonstrated that Fragle can accurately quantify ctDNA levels with targeted gene panel sequencing, we applied the method to longitudinal targeted sequencing samples from four late-stage colorectal cancer patients. In these samples, we wanted to explore the temporal relationship between Fragle-estimated ctDNA dynamics and disease progression measured by radiographic imaging (RI). Firstly, we observed strong temporal correlations between mutation VAFs and Fragle ctDNA levels across the longitudinal samples from the four patients (Fig 5a-d; Suppl. Data 15). The first patient displayed concordant and increasing VAFs and Fragle ctDNA levels, consistent with the emergence of progressive disease (PD) via RI at late timepoints (Fig 5a). The second patient developed a partial response to FOLFOXIRI treatment, consistent with both reductions in VAFs and Fragle ctDNA levels (Fig 5b). The next two patients showed a similar disease progression trajectory via RI, with initial stable disease evolving into progressive disease following multiple rounds of treatment. ctDNA dynamics inferred by Fragle showed a consistent pattern of disease progression, with ctDNA levels remaining high at all timepoints (>10%; Fig. 5c-d). While the automated variant calling pipeline failed to detect mutations at late timepoints despite the presence of PD, manual inspection of sequencing reads at these positions confirmed the presence of TP53 and ATR mutations in these samples (4-5% VAF, Suppl. Data 16). We finally considered a metastatic colorectal cancer patient for whom we had collected 21 serial blood plasma samples over a cetuximab/chemotherapy treatment course of 3 years (Fig. 5e; Suppl. Data 15). In this patient, we observed an overall temporal correlation of Fragle-based ctDNA levels, mutation VAFs, and treatment response determined from RI. However, the dynamic range of VAFs varied extensively across different mutations and time points, highlighting the challenge in estimating absolute ctDNA levels from VAFs. For example, the patient had mutations in APC and TP53, two common clonal driver mutations in colorectal cancer. The VAFs for these two mutations differed markedly, with TP53 mutation allele frequencies more than 2-fold higher at many time points (e.g. days 779 and 834). In these samples, Fragle provided an orthogonal and independent measure of ctDNA levels. Overall, these data demonstrate high concordance to Fragle-estimated ctDNA levels and disease progression estimated from radiographic imaging. Secondly, they outline how Fragle could be used to interpret and resolve heterogeneous and variable mutation VAFs profiled with targeted sequencing assays.

**Fig. 5.**
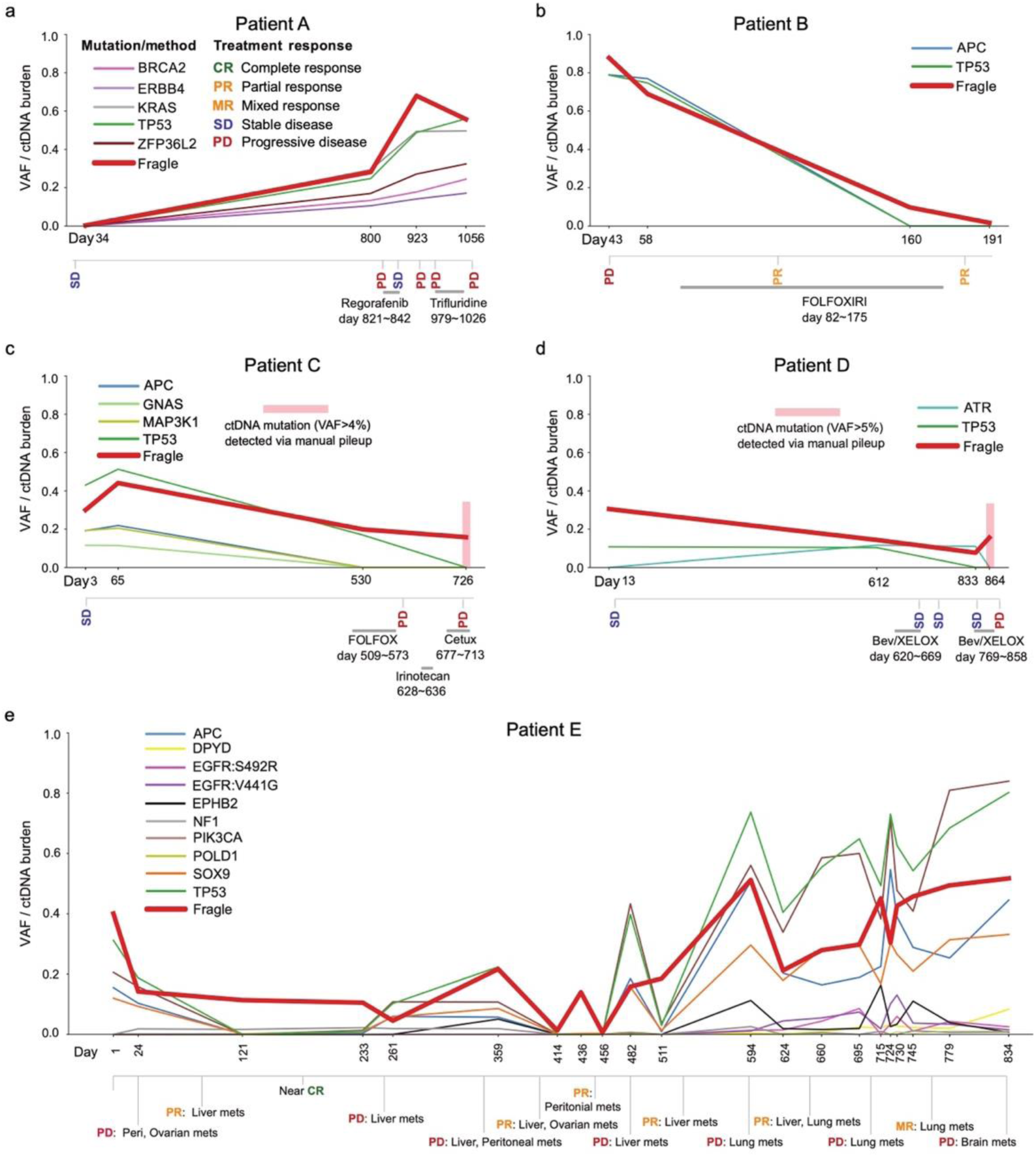
Monitoring of ctDNA levels and disease progression from targeted sequencing. **a-e**) Simultaneous longitudinal profiling of Fragle ctDNA levels and mutation VAFs in metastatic colorectal cancer patients using plasma targeted gene panel sequencing. Disease progression was captured with radiographic imaging. Only mutations detected in at least two timepoints for a given patient were included. Mutation VAFs were estimated using an automated pipeline, with manual pileup performed at highlighted timepoints where mutation detection failed.

### Risk stratification for early-stage lung cancer patients

Blood-based detection of minimal residual disease (MRD) following treatment has the potential to improve risk stratification and management strategies for cancer patients ^26, 27^. Given the ∼1% LoD for Fragle, we explored if the method could be used for tumor-naïve MRD screening, with no requirements for a matching tissue sample. We obtained targeted sequencing data from a published cohort (MEDAL) of 162 early-stage lung cancer patients that had plasma samples collected at the landmark timepoint (∼30 days following curative surgery) ^28^. In this study, plasma samples were subjected to a commercial tumor-agnostic targeted sequencing assay, and the authors classified samples into ctDNA positive (N = 4) and negative (N = 158) groups based on mutation VAFs (Fig. 6a, Suppl. Data 17). In the ctDNA-negative samples, we used Fragle to further sub-classify the samples into ctDNA-high (>1% ctDNA level, N = 101) and low (<1%, N = 57) groups. Intriguingly, despite these samples being classified as ctDNA-negative based on mutation VAFs in the targeted sequencing assay, the Fragle ctDNA-high group demonstrated significantly worse outcomes (*P* = 0.035, log-rank test) (Fig. 6b). Using a multivariate model, the association between Fragle ctDNA levels and outcomes was preserved (*P* = 0.055, Cox proportional hazard model) while controlling for known clinical prognostic variables such as tumor type and stage (Fig. 6c). Overall, these data demonstrate the potential clinical utility of Fragle as a supplement to a standard tumor-agnostic targeted sequencing assay. While Fragle was developed as a ctDNA quantification tool, these results also demonstrate that Fragle could be useful in certain settings where the detection of ctDNA is paramount, such as MRD detection and risk stratification without a matching tissue sample.

**Fig. 6.**
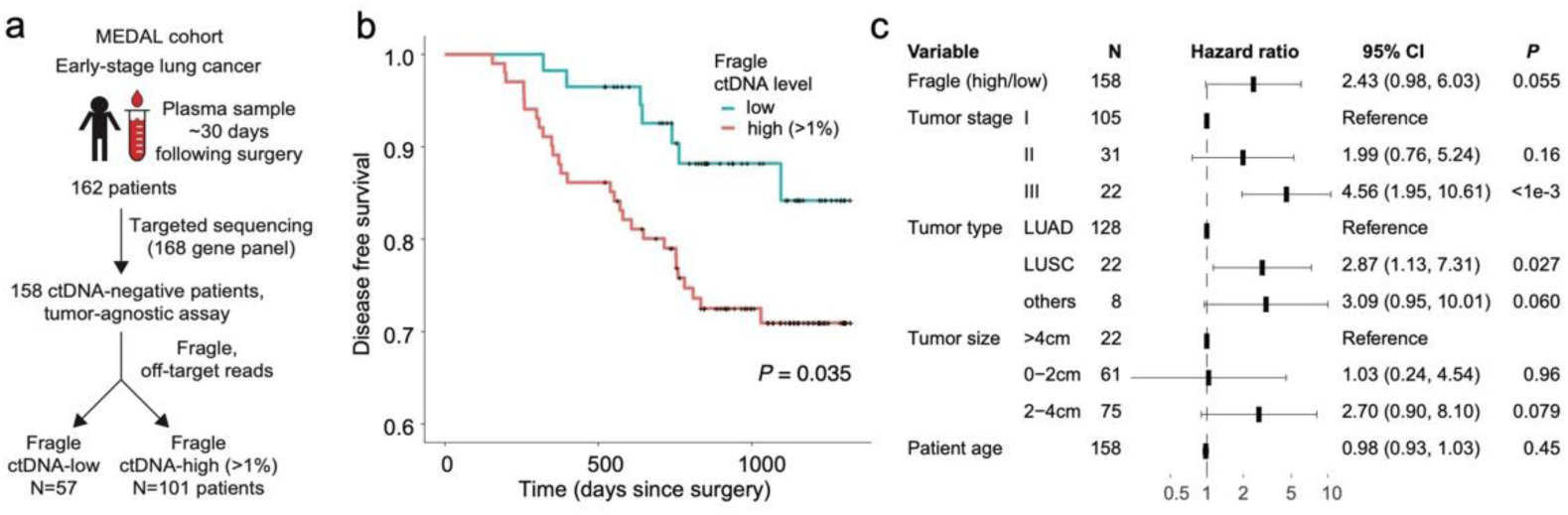
Risk-stratification of early-stage lung cancer patients: **a)** Fragle was used to predict ctDNA levels in 158 early-stage lung cancer patients classified as ctDNA-negative with a tumor-agnostic targeted gene panel assay. Plasma samples were obtained at the landmark timepoint (∼30 days after surgery) and Fragle was applied to the off-target reads to infer patients with high (>1%) and low ctDNA levels. **b)** In the 158 ctDNA-negative patients inferred with the targeted gene panel, disease free survival (DFS) was evaluated for patients with high and low Fragle ctDNA levels and compared using a log-rank test. **c)** A multivariate Cox proportional hazards model was used to evaluate the association between Fragle ctDNA levels and DFS while controlling for other clinical variables.

## Discussion

While previous studies have explored how cfDNA fragment length signatures can be used to classify plasma samples from cancer patients and healthy individuals ^16–21^, it remained unknown whether these fragmentomic signatures could also allow for accurate quantification of ctDNA levels in a blood sample. Here, we developed Fragle, a multi-stage machine learning model that quantifies ctDNA levels directly from the cfDNA fragment length density distribution, with no requirement for tumor biopsy or matched normal sample. Fragle leveraged fragmentomic features common across multiple cancer types to robustly quantify ctDNA in cancer patients, and its development and validation involved analyzing lpWGS data from 8 cancer types and targeted sequencing data from 6 cancer types. Specifically, using an in-silico data augmentation approach, we trained and evaluated Fragle on around four thousand lpWGS samples spanning multiple cancer types and healthy cohorts. Using both in vitro and in silico dilution data from unseen samples, Fragle demonstrated accurate quantification of plasma ctDNA levels with a lower LoD than the current state-of-the-art approaches for ctDNA quantification using lpWGS data. We note that Fragle has been developed and validated exclusively with whole-genome and targeted cfDNA sequencing data, further studies would be needed to evaluate if Fragle could be applied to other sequencing modalities such as bisulfite sequencing data. Moreover, modeling distinct orthogonal fragmentomic features alongside copy number profiles could unlock new opportunities to further enhance quantitative ctDNA profiling methods.

Fragle is the first method to report accurate ctDNA quantification directly from a targeted sequencing assay. Existing methods developed for ctDNA quantification are not directly compatible with targeted sequencing data, requiring either low-pass whole genome sequencing (lpWGS) data ^7^, DNA methylation data ^8, 9^, or modifications to the targeted sequencing panel ^10^. Using colon, breast, and gastric cancer plasma samples sequenced with both lpWGS and targeted gene panels, we demonstrate high concordance of Fragle estimates across assays. Furthermore, we demonstrated increased accuracy when input data was limited to the off-target reads from the targeted assay. Interestingly, while off-target reads are often filtered and ignored in a targeted sequencing experiment, these reads generally spread across the whole genome potentially mimicking ultra-lpWGS data ^29^. We used this feature to analyze longitudinal targeted sequencing samples from colorectal cancer patients, demonstrating strong concordance of Fragle-inferred ctDNA dynamics and tumor progression measured from radiographic imaging. This analysis also highlighted patients where the dynamic range of mutation VAFs varied extensively across different mutations and time points. Under these conditions, ctDNA quantification using Fragle could provide an orthogonal approach to interpret and resolve heterogeneous mutation VAFs profiled with targeted sequencing.

We also explored the potential for detecting MRD with Fragle. In a cohort of early-stage lung cancer patients with MRD evaluated at the landmark timepoint following surgery, ctDNA levels estimated by Fragle could risk-stratify patients that had otherwise been classified as ctDNA-negative using a commercial tumor-agnostic targeted sequencing assay. This result highlights the potential clinical utility of Fragle for MRD classification in settings where tumor-informed sequencing assays are not feasible or available. While tumor-informed ctDNA detection approaches offer increased MRD detection sensitivity and accuracy^28, 30^, these methods impose additional requirements for tissue sample availability, sequencing, computing, and logistics. In contrast, a tumor-naïve MRD classification approach could be applied directly to a plasma sample. Our analysis demonstrates how Fragle has potential to enhance the baseline risk stratification provided by a standard tumor-naïve targeted sequencing panel.

Fragle showed robust performance across plasma samples from 10 solid tumor types and distinct healthy cohorts. We observed strong concordance with radiographic imaging and tumor VAFs in longitudinal samples from colorectal cancer patients undergoing targeted and cytotoxic therapy. These results suggest that the machine learning approach was able to learn properties of ctDNA fragmentation that generalize across cancer types and distinct therapeutic challenges. Since Fragle uses off-target reads to quantify ctDNA with targeted sequencing, we expect the method to generalize across distinct targeted sequencing panels. While we evaluated the method using multiple targeted gene panels, future studies are needed to further characterize the performance using additional gene panels, unseen tumor types, and therapeutic exposures. Clonal hematopoiesis of indeterminate potential (CHIP) is a known contributor of cfDNA fragments in some patients, with CHIP mutations reported to occur at ∼1-2% VAFs ^24^. While we demonstrated a ∼1% LoD using in vitro and in silico diluted plasma samples, further evaluation in subjects with confirmed high levels of CHIP is needed to determine if Fragle can robustly discriminate between fragmentomic signatures from CHIP and solid tumor cells.

Fragle is fast and flexible, estimating ctDNA levels in less than a minute using paired-end cfDNA profiling and without the need for a matching tumor or buffy coat sample. By also enabling orthogonal ctDNA quantification from targeted sequencing assays, the method could limit the need for running multiple assays for disease monitoring and interpretation of negative results from plasma genotyping ^31^. This could enable simultaneous discovery of actionable cancer mutations and accurate estimation of ctDNA levels with a single assay. Overall, Fragle is a versatile and accurate method for profiling of ctDNA dynamics with potential for broad clinical utility.

## Materials and Methods

### Plasma sample collection and processing

The discovery cohort was composed of WGS plasma samples obtained from internal cohorts as well as from previous studies ^10, 12, 16, 32^. Similarly, the test cohort was composed of internal samples as well as samples from a previous study ^32^, all described in Suppl. Data 1. For new samples generated as part of this study, volunteers were recruited at the National Cancer Centre Singapore, under studies 2018/2709, 2018/2795, 2018/3046, 2019/2401, and 2012/733/B approved by the Singhealth Centralised Institutional Review Board, as well as for volunteers recruited from National University Health System (NUHS). Written informed consent was obtained from patients. Clinical data for the patients included in this study has been listed in Suppl. Data 18. Plasma was separated from blood within 2 hours of venipuncture via centrifugation at 10 min × 300 g and 10 min × 9730 g, and then stored at −80 °C. DNA was extracted from plasma using the QIAamp Circulating Nucleic Acid Kit following the manufacturer’s instructions. Sequencing libraries were made using the KAPA HyperPrep kit (Kapa Biosystems, now Roche) following the manufacturer’s instructions and sequenced on Illumina NovaSeq6000 system. Low-pass WGS (∼4x, 2×151 bp) was performed on cfDNA samples from cancer patients and healthy individuals. We used bwa-mem ^33^ to align WGS reads to the hg19 human reference genome.

### Estimation of ctDNA fractions in the discovery cohort

We estimated the ctDNA fractions in the plasma samples of 4 cancer types using distinct orthogonal methods. 12 CRC and 10 BRCA plasma samples had ∼90x cfDNA and ∼30x matched buffy coat WGS data, and their ctDNA fractions were estimated using four tumor tissue-based methods ^34–37^ as previously reported ^10^. 53 CRC samples had lp-WGS and targeted nucleosome-depleted-region sequencing data, and we inferred ctDNA fractions by averaging ichorCNA and NDRquant estimates ^10^ in these samples. The remaining 55 CRC and 57 BRCA samples only had lpWGS data and their ctDNA fractions were inferred using ichorCNA. The liver (HCC) and the ovarian cancer (OV) datasets only had lpWGS data and ichorCNA ^7^ was used to quantify ctDNA levels in these samples. The details of the estimation of ctDNA fractions are provided in Suppl. Data 2.

### Discovery cohort data augmentation approach

After identifying the plasma samples with ctDNA level ≥3% and with at least 10 million fragments in the discovery cohort, we split the cancer samples (n = 164) into training and validation sets. We repeated this 10 times, creating 10 training-validation set pairs. We then diluted each cancer plasma sample with reads from a random control plasma sample to generate in silico spike-ins, followed by down-sampling to 10 million cfDNA fragments per sample. We generated in silico samples with variable ctDNA fractions ranging from 10^-6^ up to the undiluted fractions (Suppl. Data 3). To minimize information leakage to the validation set, we evenly split the healthy control samples (n = 101) into two sets. These two control sets were then used to dilute cancer samples in the training and validation sets, respectively (see Suppl. Fig. 1).

### Overview of Fragle

Fragle quantifies ctDNA levels from a cfDNA fragment length histogram. Using paired-end sequencing data, we computed the length of each sequenced cfDNA fragment, excluding duplicates and supplementary alignments and only keeping paired reads mapping to the same chromosome with a minimal mapping quality of 30. The machine learning model consists of two stages (see Fig. 1 and Suppl. Fig. 12 for details): a quantification and a model-selection stage. The quantification stage employs two sub-models: (i) the low ctDNA burden sub-model, and (ii) the high ctDNA burden sub-model. These sub-models were designed and optimized to quantify accurately in low (<3%) and high ctDNA fraction (≥3%) samples, respectively. In the initial stage, for any given cfDNA sample, we individually input its processed fragment size profile into the two parallel sub-models. These two parallel sub-models, each with distinct loss functions, focus on ctDNA quantification for low- and high-ctDNA samples, respectively. The two parallel sub-models independently output their estimated ctDNA fractions. In the second stage, an SVM model selects the final predicted fraction from these two independent estimates. To train a final Fragle model based on the discovery cohort, we first trained a Fragle model on the training samples to obtain their ctDNA burden estimates. Among samples with ichorCNA-only ground truth ctDNA estimates, we excluded samples with a large deviation of ctDNA fractions estimated by ichorCNA and Fragle (i.e. relative difference >50% for samples with ctDNA fraction >20% by ichorCNA, >40% for samples with ctDNA fraction of 10-20%, and >30% for samples with ctDNA fraction of 3-10%). The final Fragle model was subsequently trained using all remaining samples in the discovery cohort. Notably, no samples were filtered out or selected from the unseen cohorts used to validate the final Fragle. As a result, Fragle remains entirely independent from these unseen cohorts (Fig. 1).

### Fragle model feature extraction

The feature extraction steps have been illustrated in Suppl. Fig 12. We computed the fragment length profile for all fragments sized 51-400 bp using paired-end reads with Pysam ^38^. The length profile of each sample was normalized using the highest observed fragment length count, followed by log_10_ scaling of these sample-wise normalized counts. Next, a moving average normalization (z-score of 32-nt window) was performed sample-wise for smoothing.

The transformed length features for a given sample were further standardized relative to the training set. To explore the fragmentomic feature space predictive of cancer, we identified fragment length intervals that differed between cancer samples and healthy individuals in the discovery cohort (Suppl. Fig. 13, Suppl. Data 19). The most predictive length intervals comprised both short and long cfDNA fragments, including 125-140 bp, 170-208 bp, and 246-306 bp (*P* < 10^-20^, Wilcoxon rank sum test). 281 out of 350 fragment lengths showed significant differences between cancer and healthy samples (*P* < 0.01), these were selected as candidate length features for model development.

### High and low ctDNA burden sub-model architecture

The model was implemented as a neural network with a feature embedding layer, 16 fully connected layers with batch normalization ^39^ and residual connections ^40^ (Suppl. Fig. 12). Dropout regularization ^41^ was used in between intermediate layers with a dropout rate of 30% to minimize model overfitting. A composite loss function was used combining a quantification loss and a binary cross entropy loss. The main differentiating factor between the low and the high-ctDNA burden sub-models was the loss function. Here, the high ctDNA sub-model utilizes mean absolute error (MAE) loss, while the low ctDNA burden model uses relative MAE loss:

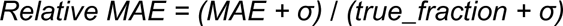

“Relative MAE” is more sensitive to prediction errors in low-ctDNA samples, because these samples have smaller ‘true_fraction’ values in the denominator. As a result, the relative MAE tends to be larger for such cases. During model optimization, this loss function encourages the model to also focus on the prediction accuracy of low-ctDNA and healthy samples, aiming to minimize the overall relative MAE. Here, σ is a hyperparameter tuned based on cross-validation data.

### Selection model

The low and high ctDNA sub-models individually predict ctDNA fractions for each sample. These two predictions are used as input features for the selection model. The training sample ground truth is labeled as 0 or 1 when the expected ctDNA fraction is <3% or ≥3%, respectively. The selection model is a binary support vector machine classifier with radial basis kernel function.

### ichorCNA and 4-feature model benchmarking

We utilized ichorCNA according to its usage guidelines, employing the default parameters to compute read count coverage with the HMMcopy Suite, followed by deducing tumor fractions with the ichorCNA R package. For the 4-feature model, we extracted four features from the fragment length profile according to a previously published study ^16^: 10-bp amplitude, and proportions of fragments sized 160-180 bp, 180-220 bp, and 250-320 bp. We used these four features to develop a random forest regression model for estimating ctDNA fractions.

### Targeted sequencing assay

Plasma and patient-matched buffy coat samples were isolated from whole blood within two hours from collection and were stored at −80 °C. DNA was extracted with the QIAamp Circulating Nucleic Acid Kit, followed by library preparation using the KAPA HyperPrep kit. All libraries were tagged with custom dual indexes containing a random 8-mer unique molecular identifier. Targeted capture was performed on the plasma samples in the unseen colorectal cancer dataset (N = 109) and in the unseen metastatic gastric cancer (N = 49) dataset, using an xGen custom panel (Integrated DNA Technologies) of 225 cancer driver genes. We also performed targeted sequencing of six plasma samples from healthy individuals to identify and blacklist unreliable variants likely attributed to sub-optimal probe design. Paired-end sequencing (2 × 151 bp) was done on an Illumina NovaSeq6000 system.

### Variant calling

FASTQ files generated from targeted sequencing were pre-processed to append unique molecular identifiers (UMIs) into the fastq headers, followed by read alignment using bwa-mem ^33^. We then performed UMI-aware deduplication using the fgbio package (https://github.com/fulcrumgenomics/fgbio). We grouped reads with the same UMI, allowing for one base mismatch between UMIs, and generated consensus sequences by discarding groups of reads with single members. To identify single-nucleotide variants and small insertions/deletions in the cfDNA samples, we first performed variant screening using VarDict ^42^ using a minimal VAF threshold of 0.05%, and annotated all variants using Variant Effect Predictor ^43^. We removed low-impact variants such as synonymous variants, and low-quality variants such as those that fail to fulfill the minimum requirements of variant coverage, signal-to-noise ratio, and number of reads supporting alternative alleles. Finally, we removed population SNPs found in Genome Aggregation Database (gnomAD) and 1000 Genomes. To further minimize false positive variants, we used duplexCaller ^44^ to identify variants with double-strand support and discarded blacklisted variants that were recurrently found in the plasma of two or more healthy individuals. Finally, when available, we identified high-confidence variants by taking advantage of serial plasma samples collected from the same patient, keeping only variants that were detectable in at least two serial samples, with VAF more than 3% in at least one sample.

### Application of Fragle to targeted sequencing data

Duplicates were removed from the targeted sequencing data using Picard MarkDuplicates function (https://broadinstitute.github.io/picard/), and on-target and off-target reads were extracted from the BAM files using samtools view function. The resulting reads were used to generate the input fragment length histograms as detailed above. We obtained targeted sequencing data of the plasma samples in the unseen colorectal cancer dataset (N = 109) and in the unseen metastatic gastric cancer (N = 49) dataset, based on a panel of 225 cancer driver genes, as described above. The targeted sequencing data for the colorectal and breast cancer datasets in the discovery cohort have been reported in the previous studies ^10, 45^, based on a panel of 100 genes of colorectal cancer mutation and a panel of 77 genes of breast cancer mutation, respectively. Summary statistics for all gene panels such as gene count, genomic coverage, target regions, and on-target coverage ratio have been provided in the supplemental material (Suppl. Data 20-22). In an additional analysis, we used targeted sequencing data from 116 patients profiled with the Foundation Medicine Liquid CDx assay. Samples belonged to different cancer types such as lung, colon, breast, pancreas, and uterus cancer (Suppl. Data 14). We filtered known germline variants using gnomAD (v4) and analyzed variant allele frequencies using all remaining variants reported by the company. Since Fragle requires off-target BAM files for prediction, we constructed a targeted sequencing bed file using the 311 genes reported to comprise this panel (Suppl. Data 21).

### Lung cancer survival analysis

Plasma targeted sequencing data from the MEDAL cohort (Project ID: OEP004204) ^46^ was retrieved from National Omics Data Encyclopedia (NODE). Alignment to the human genome (hg19) was conducted using bwa-mem ^33^. Duplicates in the aligned data were marked using Samblaster ^47^. Putative target regions were identified by calculating the median coverage per base from a subset of randomly selected BAM files (n = 38). Coverage of regions without any reads was reported as zero. Next, the resulting consensus bedgraph file was segmented into 100 bp bins. Bins with median coverage exceeding 2x were selected and merged if they were within 100 bp of each other, to form contiguous regions. The resulting BED file was used for obtaining the off-target BAM file for each sample using samtools.

### Unseen in vitro dilution experiments

The first in vitro dilution experiment included high ctDNA burden cfDNA samples from 2 individual CRC patients which were selected to create a starting point for the dilution series. The ctDNA fraction for each sample was determined by 2 methods (ichorCNA ^7^ and NDRquant ^10^), with high concordance across methods (sample 1: ctDNA content 38% by ichorCNA, 37% by NDRquant; sample 2: 39% and 38%, respectively). Commercial pooled cfDNA from healthy volunteers (0% ctDNA) was purchased from PlasmaLab (lot numbers 2001011, 210302) and was used to set up a 9-point serial dilution of the ctDNA fraction for each sample (Suppl. Data 12), with 3 technical replicates per dilution point. The second in vitro dilution experiment started from 3 plasma samples of gastric cancer, that had concordant ctDNA estimates between methods (sample 1: ctDNA content 7.6% by ichorCNA, 8.7% by Fragle, sample 2: 17.2% and 14.7%, sample 3: 13.4% and 19.1%, respectively). 12 control plasma cfDNA samples were purchased from Ripple Biosolutions and were used to set up a 7-point serial dilution of the ctDNA fraction for each cancer sample (Suppl. Data 13), with 3 technical replicates. We randomly selected 3 control samples and pooled them before diluting the plasma of cancer. lp-WGS was performed with a depth of ∼ 4-5x.

### Unseen in silico dilution experiments

Our unseen in silico dilution experiment included 7 unseen breast and 13 unseen colon cancer samples each containing high and concordant ctDNA fraction estimates based on Fragle and ichorCNA (>10% ctDNA based on both methods with a relative difference < 5%). We prepared 20 healthy mixtures, each created by pooling 3 random samples from an unseen control cohort. A 6-point serial dilution for each sample was set up using these healthy mixtures to dilute the 20 cancer samples, with 20 technical replicates. A total of 2400 dilution samples were created ranging from 5% to as low as ∼0.1% ctDNA fraction, each dilution point containing 400 samples (Suppl. Data 11).

## Data availability

Published data used in this study and their access codes are present in Suppl. Data 1. Data generated in this study have been deposited at the European Genome-phenome Archive (EGA; Dataset ID: EGAD50000000167). Data are available under restricted access and will be released subject to a data transfer agreement.

## Code availability

The Fragle software is attached as Suppl. Data 23 and will be made publicly available via GitHub. The software can be directly applied to lpWGS/off-target BAM files aligned to hg19 / GRCh37 / hg38 reference genomes without any preprocessing.

## Supplementary materials

Supplementary Fig. 1-13

Supplementary Data 1-23

## Disclosure

The authors declare no competing interests.

## Acknowledgments

This work was supported by Agency for Science, Technology and Research under its IAF-PP programme (grant ID: H1801a0019) and the Singapore Ministry of Health’s National Medical Research Council under its OF-IRG program (OFIRG21nov-0083). This work makes use of data from the Chinese University of Hong Kong (CUHK) Circulating Nucleic Acids Research Group as reported previously (doi:10.1073/pnas.1500076112 and doi: 10.1158/2159-8290.CD-19-0622), and data from CRUK_CI, University of Cambridge, Rosenfeld Lab, as reported previously (doi:10.1126/scitranslmed.aat4921). The authors gratefully acknowledge Dr. Dennis Lo and his research group at CUHK, as well as Dr. Nitzan Rosenfeld and his research group at University of Cambridge, for providing access to cfDNA cohorts.

## Notes

### Competing Interest Statement

The authors have declared no competing interest.

### Summary of Updates

Figure 2 has been updated; some mistakes were there. Also experiments related to cancer stage stratification have been added. Text has been improved. New supplementary figures added.

