## Supplementary Figures for "Fragle: Universal ctDNA quantification using deep learning of fragmentomic profiles"

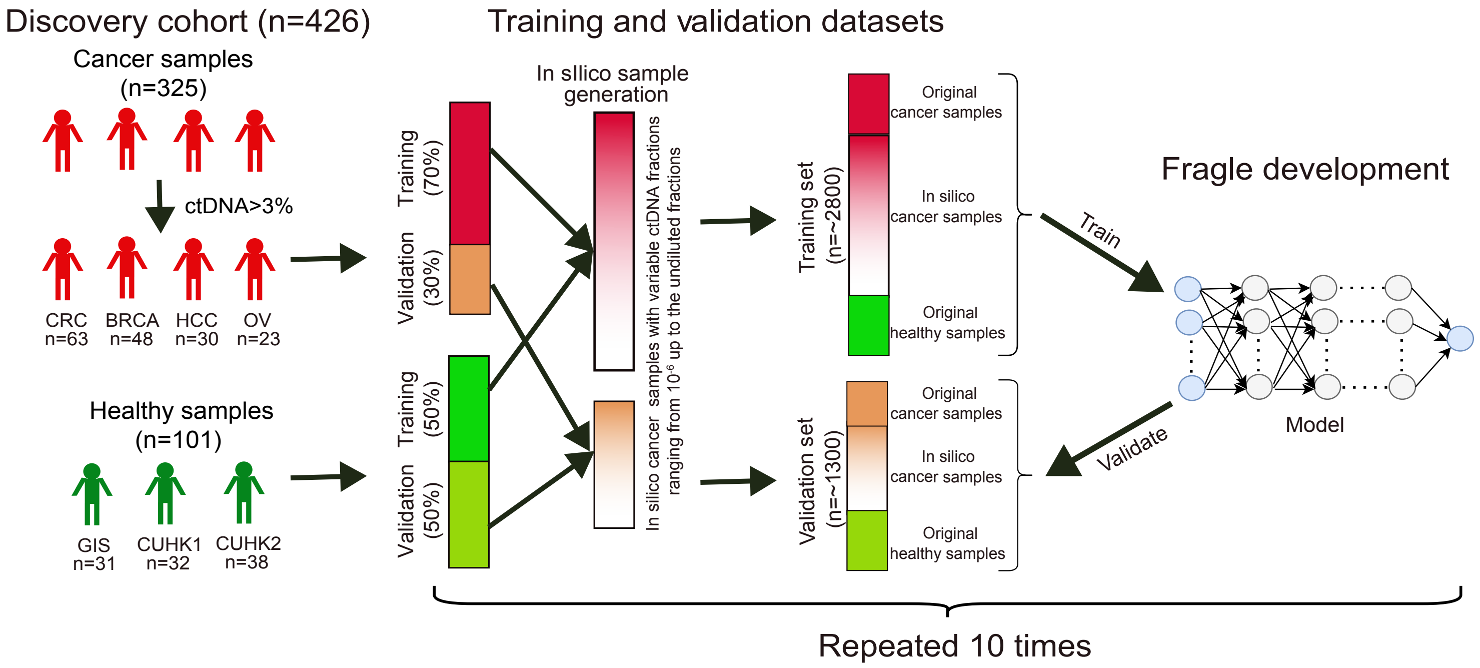


**Supplementary Fig. 1** Schematic showing how the predictive models were trained and validated using the healthy control samples, original cancer samples, and in silico cancer spike-ins in the discovery cohort.

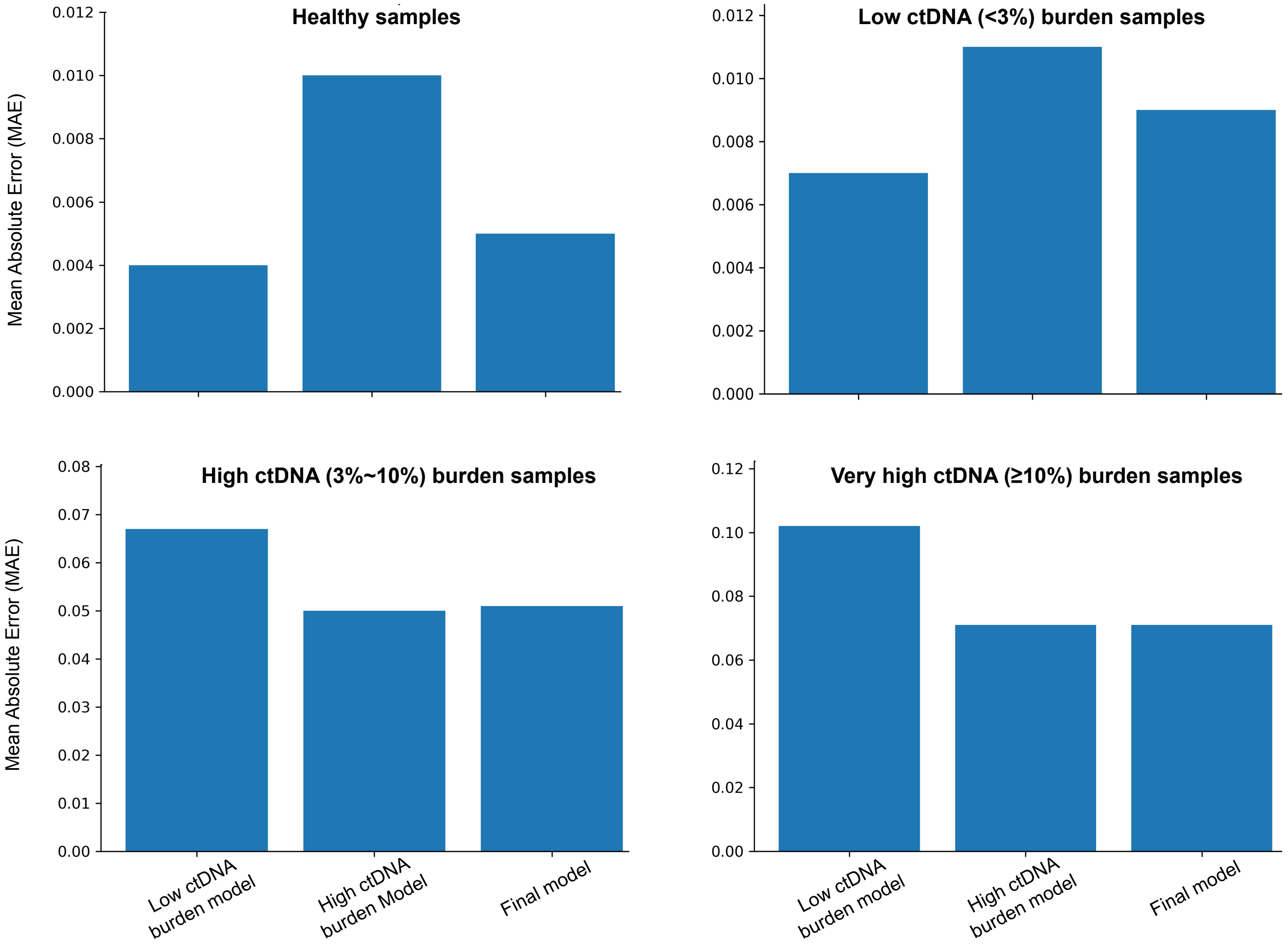


**Supplementary Fig. 2** Mean absolute error (MAE) between predicted and expected ctDNA fractions for low and high ctDNA burden models as well as the final model. Performances of the three models were compared on the basis of different sample groups that included the samples with different ctDNA levels.

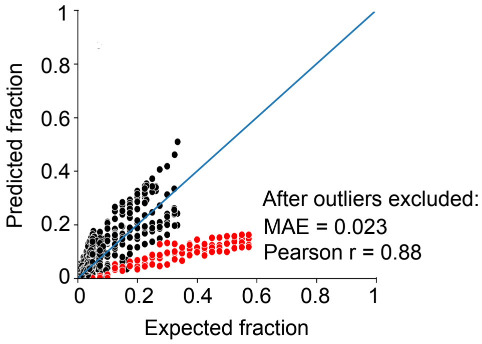


**Supplementary Fig. 3** Comparison between expected and predicted ctDNA fractions of the plasma samples of OV in the validation set. The outlier samples (red) generated from one original OV sample were excluded, improving Pearson correlation to 0.88 and MAE to 0.023.

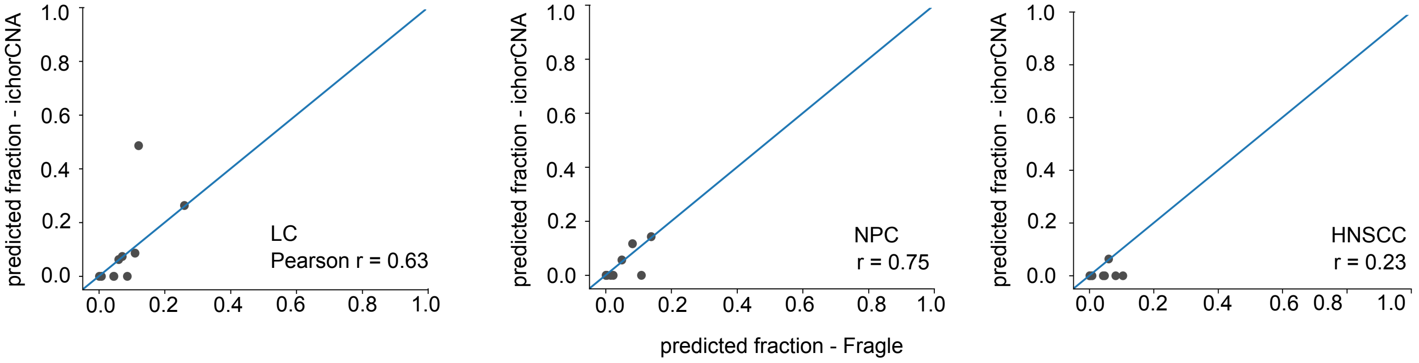


**Supplementary Fig. 4** Comparison between Fragle and ichorCNA in their predicted ctDNA fractions in unseen test samples from LC, NPC, and HNSCC patients.

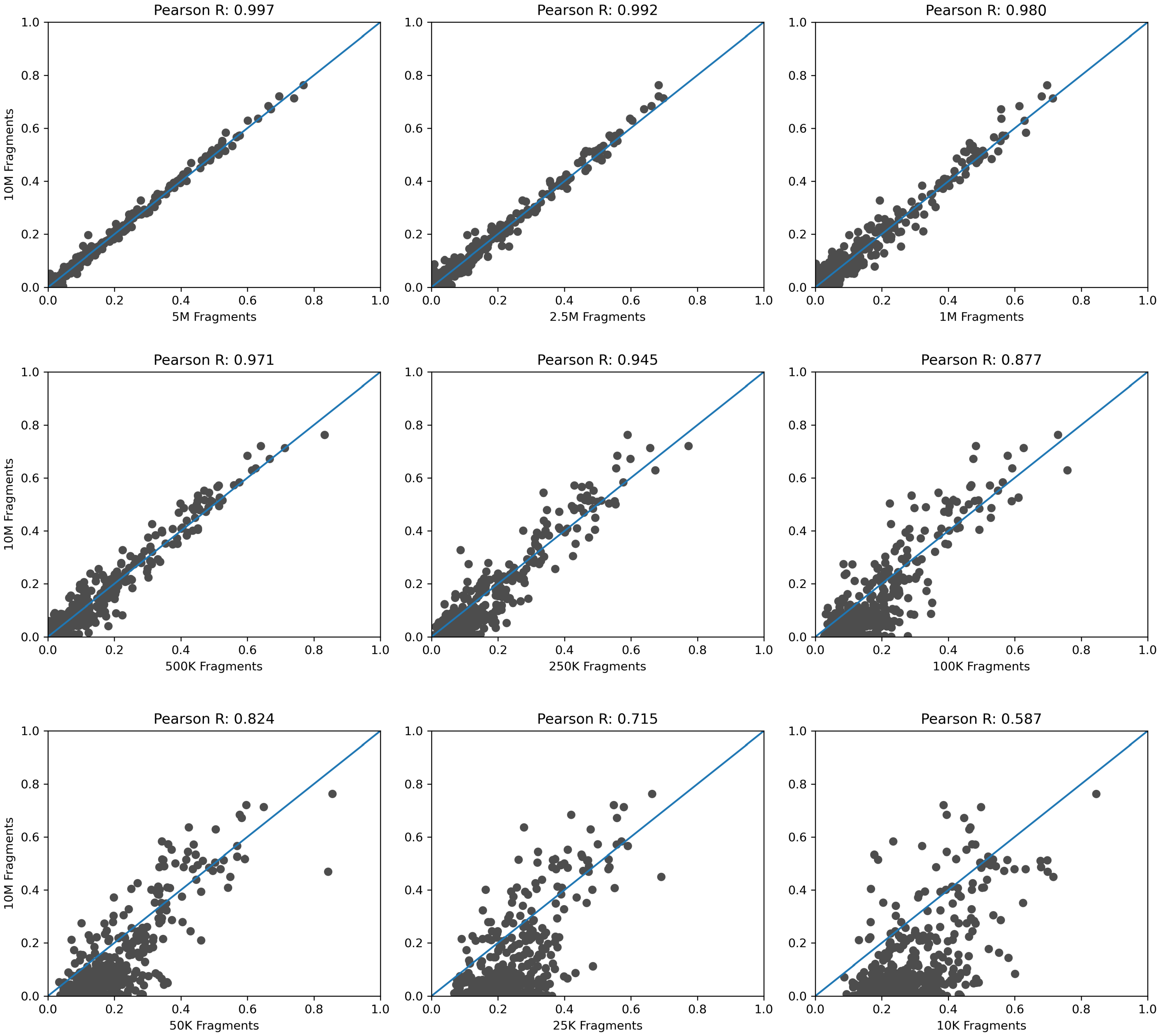


**Supplementary Fig. 5** Comparison of the Fragle-predicted ctDNA fractions in the unseen cfDNA WGS samples with 10 million cfDNA fragments and their downsampled counterparts.

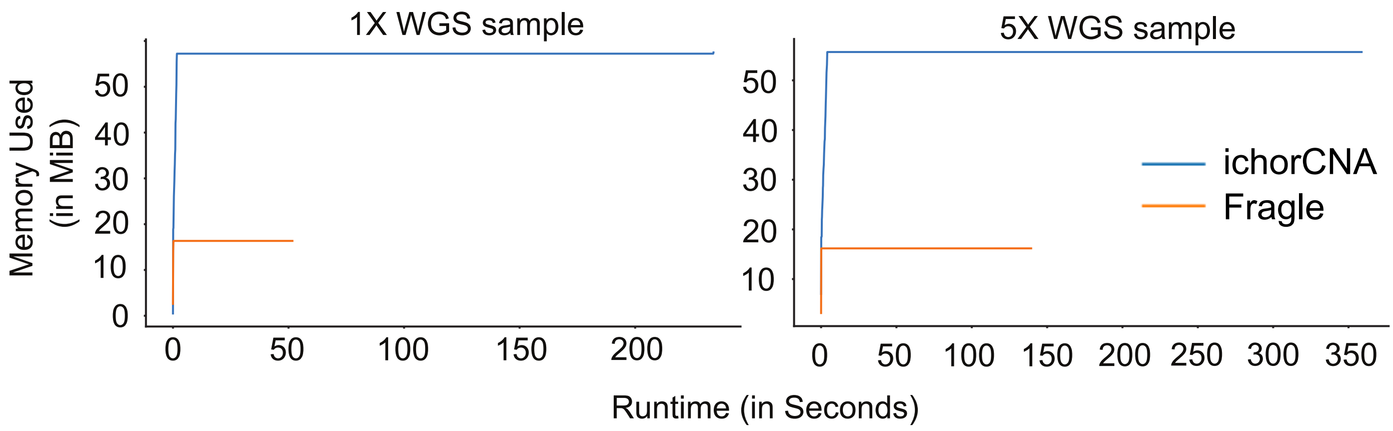


**Supplementary Fig. 6** Runtime and memory comsumption comparison between Fragle and ichorCNA for 1X and 5X WGS sample.

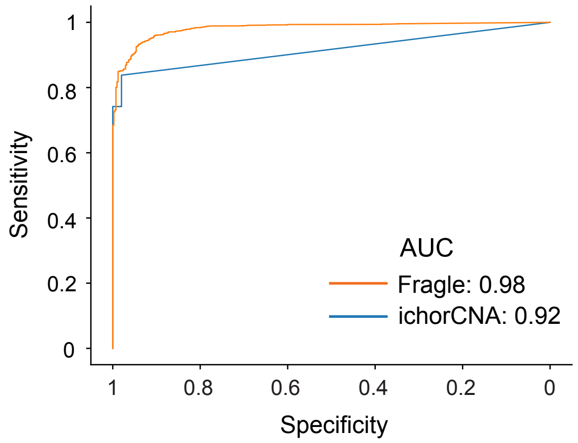


**Supplementary Fig. 7** ROC analyses for classification of healthy control and cancer samples in which ground-true ctDNA fractions were based on multiple methods.

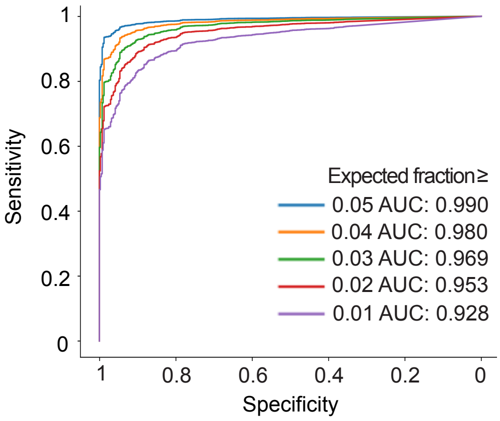


**Supplementary Fig. 8** ROC analysis to examine the diagnostic performance on the validation samples with different thresholds to exclude low ctDNA samples.

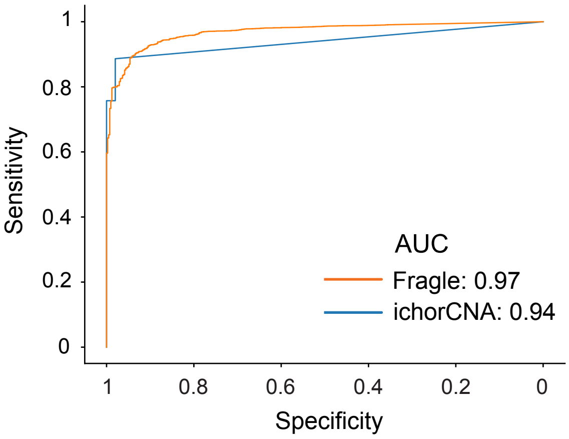


**Supplementary Fig. 9** ROC analyses for classification of healthy control and cancer (≥3% ctDNA) samples (validation samples).

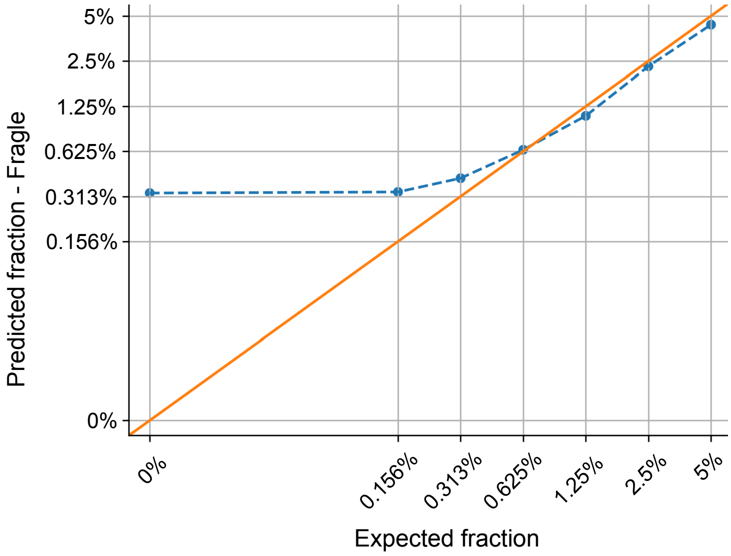


**Supplementary Fig. 10** Median ctDNA fractions based on Fragle for the in silico dilution samples with low ctDNA fractions.

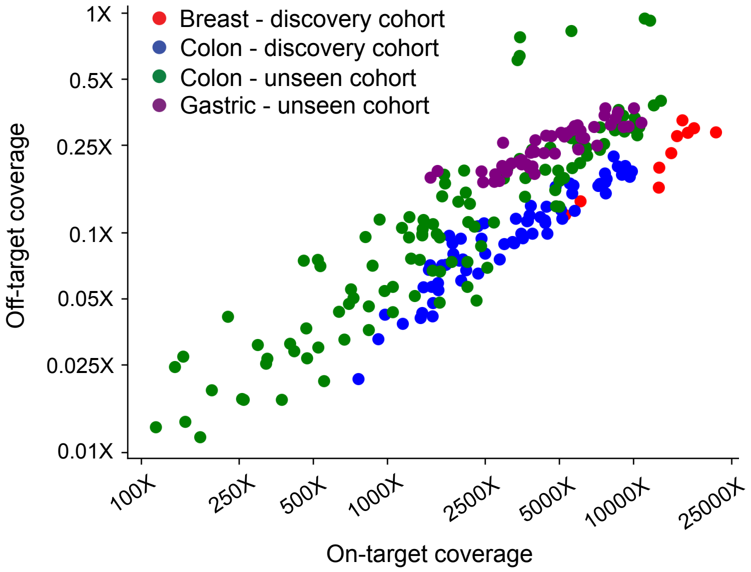


**Supplementary Fig. 11** On-target and off-target coverage extracted from the targeted sequencing data of the cfDNA samples from four different cohorts consisting of three cancer types.

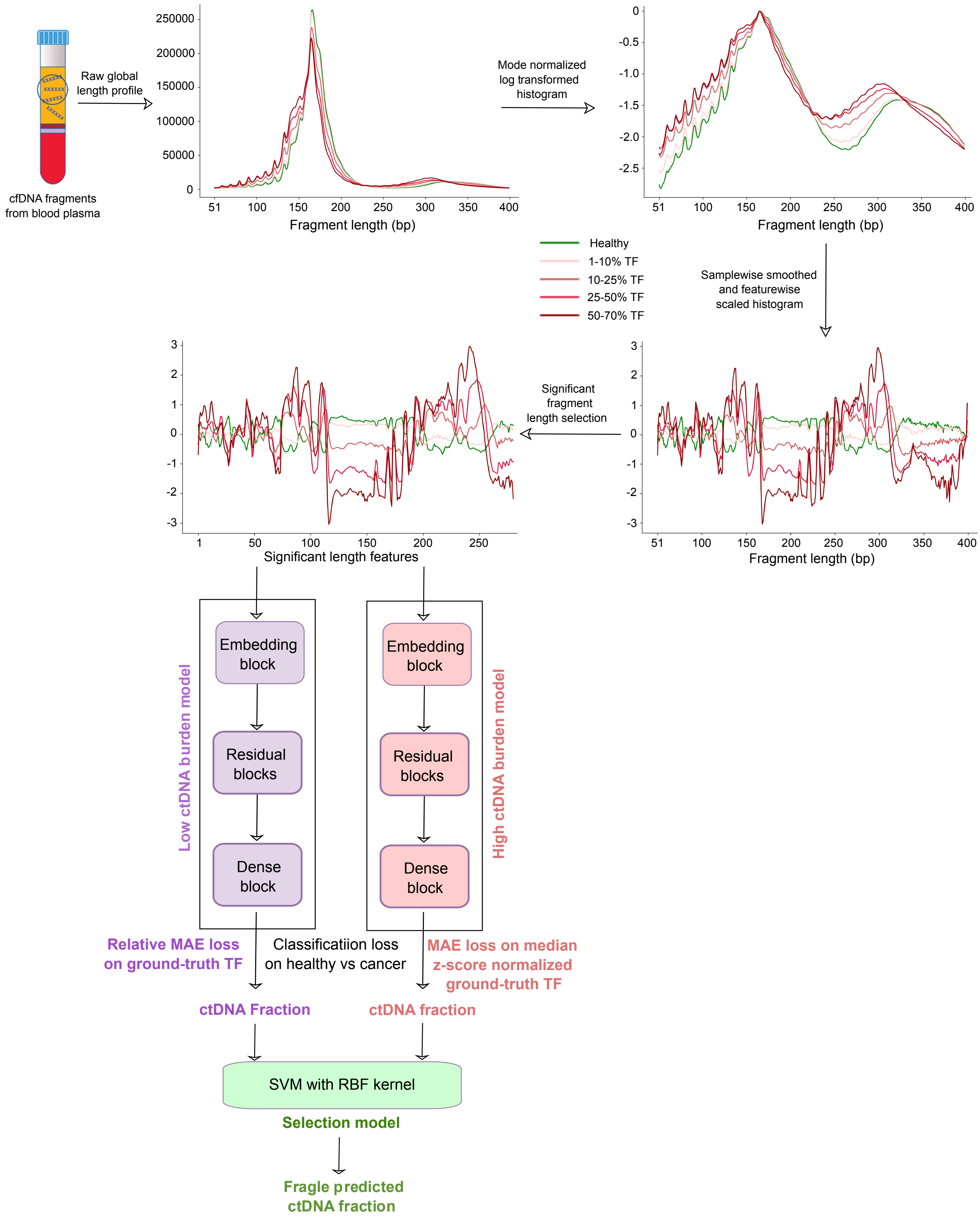


**Supplementary Fig. 12** Details about the feature processing and model structure of Fragle.

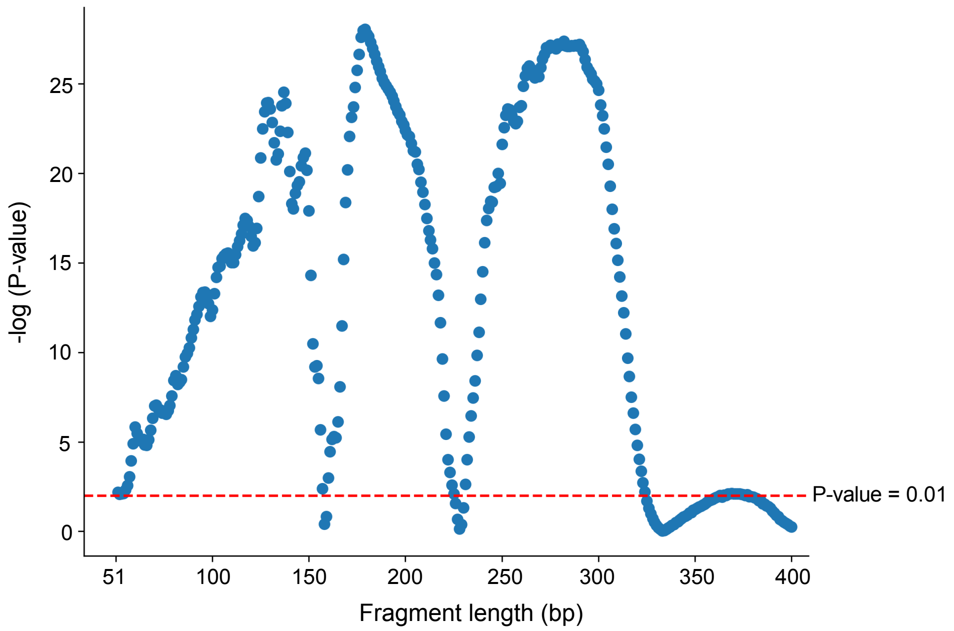


**Supplementary Fig. 13** Significance values for fragment lengths informative of distinguishing between cancer and healthy samples.
